# Human cell adaptation of the swine acute diarrhea syndrome coronavirus spike protein

**DOI:** 10.1101/2024.07.29.605615

**Authors:** Clàudia Soriano-Tordera, Rafael Sanjuán, Jérémy Dufloo

## Abstract

Swine acute diarrhea syndrome coronavirus (SADS-CoV) is a recently identified highly pathogenic swine coronavirus. In vitro, SADS-CoV can infect cell lines from many different species, including humans, highlighting its high zoonotic potential. Coronavirus spike glycoproteins play a critical role in viral entry and are involved in determining viral host range and cellular tropism. Here, we used experimental evolution to investigate how the SADS-CoV spike protein adapts to human cells and to identify potential variants with increased infectivity. We evolved a recombinant vesicular stomatitis virus expressing the SADS-CoV spike (rVSV-SADS) in three human cell lines. After ten passages, increased viral replication was observed, and spike mutations were identified by sequencing. Mutations were functionally characterized in terms of viral fitness, spike processing and fusogenicity. Our results thus identify potential human-adaptive mutations in the SADS-CoV spike that may further enhance its zoonotic potential.

**Importance:** Coronavirus transmission from animals represents a serious threat to humans. Pigs are of particular concern because of their proximity to humans and the several coronaviruses they harbor. In particular, the swine acute diarrhea syndrome coronavirus (SADS-CoV) is a recently identified highly pathogenic porcine coronavirus that has a very broad tropism in vitro, highlighting its high zoonotic risk. The coronavirus spike protein is a strong determinant of species tropism, and spike mutations may facilitate cross-species transmission. Here, to identify potential variants with increased ability to enter human cells, we used an experimental evolution approach to study how the SADS-CoV spike adapts to different human cell lines. These mutations, should they occur in nature, could potentially increase the zoonotic potential of SADS-CoV.

## Introduction

Coronaviruses can infect humans, other mammals and birds, causing respiratory and gastrointestinal disease (1). To date, seven human coronaviruses have been identified. Four of them (NL63, 229E, OC43, and HKU1) cause cold-like symptoms and three (SARS-CoV, SARS-CoV-2, and MERS-CoV) cause more severe respiratory diseases (2). Current data suggest that these viruses were transmitted relatively recently to humans from different mammals, including bats, rodents, and other intermediate animals such as civets or camels (3, 4). The transmission of novel coronaviruses from animals to humans therefore represents a significant threat to human health and highlights the importance of identifying and characterizing coronaviruses at risk of zoonosis (5, 6).

Swine acute diarrhea syndrome coronavirus (SADS-CoV) is porcine alphacoronavirus first identified in 2016 in Guangdong, China. SADS-CoV infection is associated with acute diarrhea, vomiting and mortality rates of up to 90% in piglets less than five days old, making it a significant threat to the pork industry (7–9). Several studies have demonstrated that the virus has a broad species tropism in vitro, with the ability to infect cell lines derived from a wide range of animals, including bats, rodents, pigs, chickens, non-human primates and humans (10–12). SADS-CoV can also infect other species in vivo, such as neonatal mice and chicken embryos (13–15), further highlighting its potential for cross-species transmission.

The coronavirus spike (S) glycoprotein is the primary mediator of target cell attachment and entry and is involved in determining viral host range and cellular tropism (16). The mature spike is composed of two subunits: S1, which binds the host receptor, and S2, which allows the fusion of viral and cellular membranes. Both subunits are generated by post-translational cleavage of the spike precursor S0 at the S1/S2 junction, which contains a multibasic furin cleavage site (FCS) in some coronaviruses (17), including SADS-CoV. The S1 subunit contains the two most variable regions of the spike protein: the N-terminal domain (NTD), which is often responsible for the recognition of cell surface carbohydrates, and the C-terminal domain (CTD), which usually binds to the proteinaceous receptor. In contrast, the S2 subunit contains the more conserved domains of the fusion machinery, including the fusion peptide (FP), and the heptad repeats 1 and 2 (HR1/2) (18–20)

Viral glycoproteins frequently evolve to increase human infectivity. For example, the A82V mutation in the Ebola virus glycoprotein increases viral entry into human cells and was associated with the large 2013-2016 West African epidemic (21). In coronaviruses, amino acid substitutions in the spike protein have also been associated with increased human infectivity. For example, several SARS-CoV-2 variants with heavily mutated spikes have successively spread across globally (22). Some of these mutations were associated with increased replication (23), enhanced receptor affinity (24), differential spike cleavage (25, 26), use of alternative entry pathways (27), or escape from antibody-mediated neutralization (28, 29). Therefore, studying how coronavirus spikes adapt to human cells may allow the identification of variants with increased human transmissibility, which may be useful for viral surveillance and preventing potential spillover infections.

Due to its high risk of cross-species transmission, SADS-CoV is currently classified as a biosafety level (BSL) 3 virus, which complicates its laboratory use. Nevertheless, recombinant replication-competent vesicular stomatitis viruses (rVSV) represent a safe and relevant surrogate system for the study of the entry process of highly pathogenic viruses (30, 31). In these chimeric viruses, the VSV glycoprotein is replaced with a foreign receptor-binding protein, which allows to investigate its function, evolution, or antigenicity. This system offers several advantages, including BSL-2 level containment, ease of production, and high yield. In experimental evolution approaches, this also allows to avoid using the original virus for long-term passaging experiments, as this may result in the emergence of gain-of-function mutations. Previous studies have used this system to study coronavirus spike function (31, 32) and adaptation (28, 33, 34).

Here, to study how the SADS-CoV spike adapts to human cells, we generated a recombinant replication-competent vesicular stomatitis virus expressing the SADS-CoV spike (rVSV-SADS) and serially passaged it in three human cell lines. Adaptive mutations were identified and functionally characterized in terms of spike processing, incorporation, cell-cell fusogenicity, and effect on viral fitness. Our results led to the identification of several mutations in the SADS-CoV spike that may increase entry into human cells.

## Results

### Adaptation of a rVSV-SADS to human cell lines

To study the adaptation of the SADS-CoV spike to human cell lines, we first obtained a GFP-expressing rVSV modified to express the SADS-CoV spike (rVSV-SADS) instead of the VSV envelope glycoprotein. To achieve sufficient viral titers, the rVSV-SADS had to be first passaged twice into Huh-7 liver cells, known to be susceptible to SADS-CoV infection (10, 11). This initial stock was called P0. We then challenged 48 different human cell lines from the NCI-60 panel with the P0 virus. GFP signal was detected in multiple cell lines (**Supplementary Table 1**), and we selected to top-12 lines to attempt serial passages of the virus. This was successful only for

OVCAR-8 (ovary) and H23 (lung) cells. Some NCI-60 cell lines (e.g. SW-620) were initially susceptible to the P0 virus but did not support viral spread at sufficient levels to allow serial passaging, suggesting efficient viral entry but poor egress. We thus selected Huh-7, OVCAR-8 and H23 cells for experimentally evolving rVSV-SADS. A total of 10 passages were performed in triplicate (R1-R3; **Figure 1A**). The viral population went extinct in two H23 and one OVCAR-8 replicates (**Figure 1B**). Viral titers increased along passages in all the other evolution lines (Spearman correlation: P < 0.005 in all cases), reaching up to 10^5^ FFU/mL in OVCAR-8 and 10^6^ FFU/mL in Huh-7 and H23 (**Figure 1B**).

**Figure 1.**
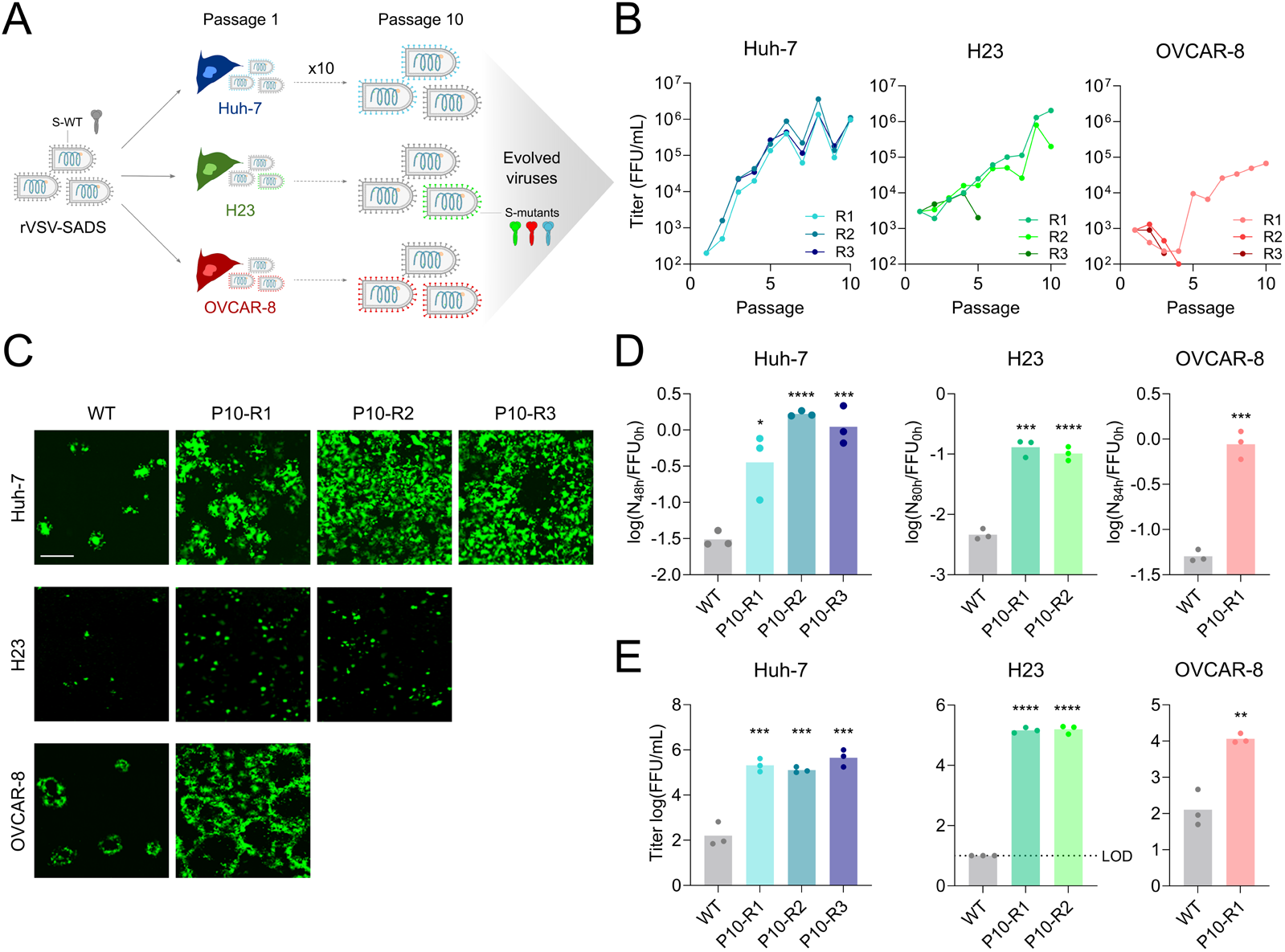
Adaptation of the rVSV-SADS to human cell lines. **A**. Schematic of the experimental evolution. rVSV expressing the SADS-CoV spike was passaged 10 times in Huh-7, OVCAR-8 and H23 cells. Three independent evolution lines (R1-R3) were performed. Evolved viral populations were characterized and sequenced at passage 10. **B**. Titration of supernatants along passages. Titers were measured after each passage as foci-forming units (FFU) per mL. The H23 R3 and OVCAR-8 R2 and R3 lineages went extinct before passage 5. **C**. Representative images of infection with the original virus (WT) and evolved lineages (passage 10) in their respective cell lines. Huh-7: 48 hpi, OVCAR-8 and H23: 72 hpi. Scale bar: 875 µm. **D**. Spread rate of WT and P10 rVSV-SADS-CoV in their respective cell lines, quantified as the GFP signal at the final time point divided by initial titer. **E**. Final titers of WT and P10 rVSV-SADS-CoV in their respective cell lines. The dotted line indicates the limit of detection. In D and E, each dot represents a technical replicate (n = 3), and an unpaired two-tailed t-test was performed comparing all evolved lineages to the WT. * P < 0.05; ** P < 0.01; *** P < 0.001; **** P < 0.0001.

To assess whether evolved lineages replicated better than the wild-type (WT) virus in human cells, each cell line was infected with the WT rVSV-SADS and the final passage (P10) of their respective evolved lineages with an approximately equal input, and two quantities were measured: the area occupied by GFP-positive cells divided by the initial titer as an indicator of viral spread in the cell cultures (spread rate), and the final titer (yield). The evolved viruses spread faster than the WT virus in the three cell lines, as shown by an >10-fold increase in the number of infected cells at endpoint in all cases (**Figure 1C-D**). Similarly, endpoint titers were between 90-fold and 15,000-fold higher than those reached by the WT (**Figure 1E**). These results suggest adaptation of the rVSV-SADS to the three human cell lines. As this could be due to mutations within the spike or elsewhere in the VSV genome, we sought to determine whether spike mutations emerged and were responsible for the observed adaptation of the rVSV-SADS to human cells.

### Identification of SADS-CoV spike mutations

To identify specific mutations within the SADS-CoV spike, we sequenced the spike gene of the six evolved viral populations at passage 10. Eight non-synonymous amino acid substitutions were identified in different functional domains of the spike (**Figure 2A** and **Supplementary Table 2**). One mutation was observed in the S1 CTD, and all the other changes occurred in S2, including inside or close to the HR1, the HR2, and the FP (**Figure 2A**). Moreover, two mutations in the S2 C-terminal domain led to the appearance of a premature stop codon, truncating 11 (Q1120*) or 1 (G1130*) amino acids from the spike cytoplasmic tail. Two of the eight mutations were present in all lineages (S1037G and T1107I). Sequencing the Huh-7-passaged founder (P0) and the original viral stock (WT) allowed us to determine that these two mutations were fixed during the two initial passages performed in Huh-7 cells to obtain P0. Nonetheless, additional mutations were observed in various P10 lineages, and some occurred in several lineages, indicating parallel evolution. For example, the V754M and G1130* mutations were observed in the two H23 lineages, and Q1120* emerged in OVCAR-8 R1 and Huh-7 R3.

**Figure 2.**
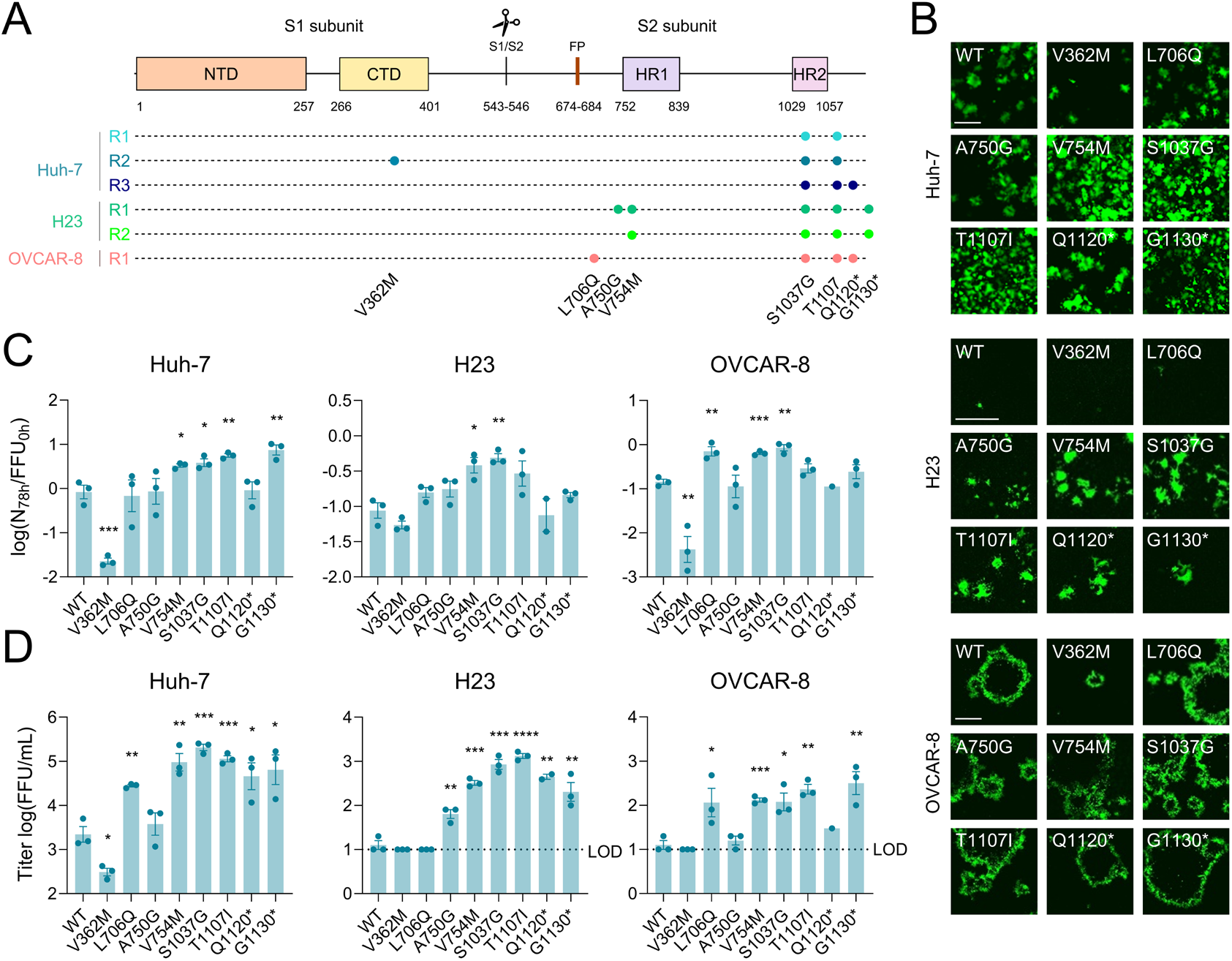
Effect of spike mutations on viral fitness components. **A**. Spike mutations in each viral lineage at passage 10. Mutations were identified by Sanger sequencing. Spike functional domains are shown as colored boxes. Each amino acid change is indicated below. * indicates the apparition of a premature stop codon. **B**. Representative images of infection with WT and mutant rVSV-SADS in each cell line. Huh-7: 48 hpi, H23 and OVCAR-8: 120 hpi. Scales bars: 800 µm. **C**. Spread rate of WT and mutant rVSV-SADS in each cell line, quantified as the GFP signal at the final time point divided by initial titer. **D**. Final titers of WT and mutant rVSV-SADS in each cell line. The dotted line indicates the limit of detection. In C and D, the bar represents the mean ± SEM and each dot represents a technical replicate (n = 1-3). An unpaired two-tailed t-test was performed comparing all mutants to the WT. * P < 0.05; ** P < 0.01; *** P < 0.001; **** P < 0.0001.

### Effect of the SADS-CoV spike mutations on viral fitness

To test the effect of all S mutations on the rVSV-SADS fitness, we used site-directed mutagenesis to generate rVSV-SADS variants carrying each point mutation individually. We infected the three cell lines with the WT and mutant viruses and measured viral spread (**Figure 2B, 2C**) and endpoint titers (**Figure 2D)**. Some mutations increased these fitness indicators in the three cell lines used. For example, the V754M and S1037G variants spread >4-fold faster than the WT virus in all cell lines (**Figure 2C**). These two mutants, as well as the T1107I and G1130* variants also reached higher titers than the WT in all cell lines (**Figure 2D**). In contrast, some mutations had a cell type-dependent effect. For example, the A750G mutant, which appeared in H23, only replicated to higher titers than the WT (5-fold) in this cell line, suggesting that it may be a cell-type specific adaptation of the SADS-CoV spike (**Figure 2D**). Similarly, the L706Q mutation, which appeared in OVCAR-8, only showed enhanced viral spread in this cell line (4.9-fold; **Figure 2C**) and replicated to higher titers in OVCAR-8 and Huh-7 (9.2- and 12.6-fold, respectively), but not in H23 (**Figure 2D**). The Q1120* mutant did not spread faster in any of the cell lines but reached higher titers than the WT in Huh-7 and H23 cells (**Figure 2B-D**). Finally, surprisingly, the V362M mutation was neutral or deleterious in all cell lines.

### Effect of mutations on SADS-CoV spike fusogenicity and processing

To gain mechanistic insights into how most of these mutations increase rVSV-SADS fitness in human cells, we first measured the fusogenicity of all mutant spikes using a GFP complementation assay (**Figure 3A**). Most spikes induced cell-cell fusion at levels comparable to the WT, except the V362M and L706Q mutations, which reduced syncytia formation by 2.6-fold and 2.3-fold, respectively (one-way ANOVA: P < 0.001 in both cases). Interestingly, the V362M mutation had a detrimental effect in all cell lines, whereas the L706Q was beneficial in OVCAR-8 and Huh-7, suggesting that there is no direct link between SADS spike cell-cell fusogenicity and infectivity in human cells (**Figure 2B-D**). Finally, using Western blot, we quantified the effect of the mutations on spike processing and incorporation into rVSV-SADS (**Figure 3B**). Incorporation and cleavage of the Q1120* and G1130* mutants could not be measured due to the premature stop codon. The V362M and, to a lower extent, the A750G and S1037G mutations decreased cleavage compared to the WT spike, showing a 6.5-, 1.8-, and 1.6-fold decrease in the S2/(S2+S0) ratio, respectively. Given the known link between SADS-CoV spike cleavage and cell-cell fusion (35), the lower spike processing of the V362M mutant likely explains its reduced cell-cell fusogenicity. Finally, we observed that all mutant spikes, except the A750G, were better incorporated into rVSV-SADS, as indicated by 1.7 to 4.2-fold increase in the (S0+S2)/VSV-M ratio, which may be an additional factor explaining the increased fitness of the mutants in human cells.

**Figure 3.**
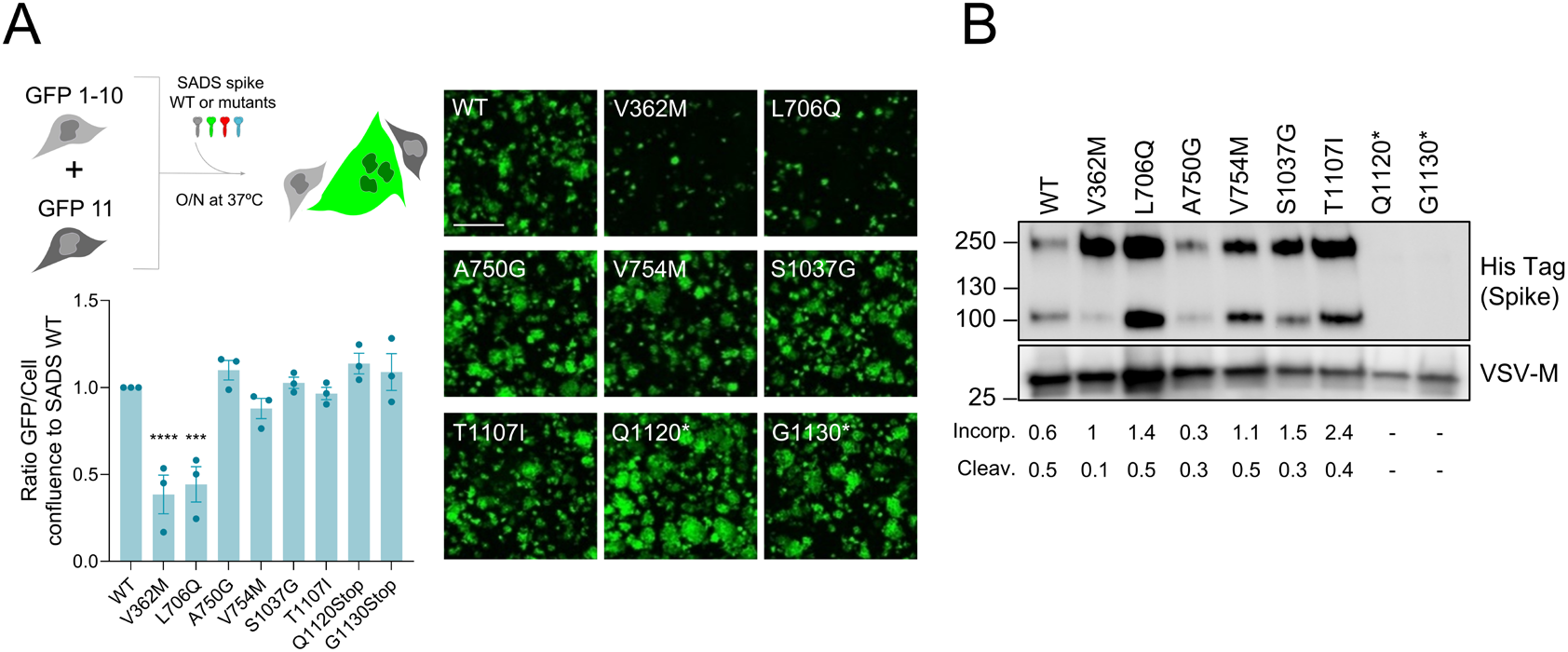
Effect of mutations on SADS-CoV spike fusogenicity and processing. **A**. Cell-cell fusogenicity of the spike mutants. Top left: schematic of the cell-cell fusion assay. Briefly, HEK293T GFP-Split cells were mixed and transfected with SADS-CoV spike and GFP-positive syncytia were quantified. Bottom left: data show as the log_10_ of the GFP confluence divided by cell confluence ratio, normalized to the SADS WT spike (mean ± SEM, n = 3 independent experiments). A one-way ANOVA with Dunnett’s correction for multiple tests was performed. *** P < 0.001; **** P < 0.0001. Right panel: representative images of spike-mediated cell-cell fusion at 14 h post-transfection. Scale bar: 400 µm. **B**. Spike Western blot of WT and mutant rVSV-SADS particles. The spike was detected with an anti-His-Tag Antibody and VSV-M was used as a loading control. The C-terminally truncated Q1120* and G1130* mutant spikes could not be detected because they do not express the C-terminal His-Tag. Spike incorporation was calculated as the proportion of total spike (S0+S2) over VSV-M protein. Spike cleavage was calculated as the proportion of S2 over total spike (S0+S2).

## Discussion

Here, we experimentally evolved the SADS-CoV spike to investigate its adaptation to human cells and to identify potential variants with increased human infectivity. To this end, we serially passaged a SADS-CoV spike-expressing rVSV in human cell lines. After 10 passages, all the evolved lineages exhibited enhanced viral replication compared to the WT virus. Moreover, eight spike mutations were identified in these evolved lineages, most of which increased rVSV-SADS spread and yield, either in all cell lines or in a cell type-specific manner. We did not determine whether these mutations were truly human-adaptive or whether they may generally increase viral fitness in any cell line susceptible to SADS-CoV entry, regardless of its species of origin. To answer this question, we tried to infect the porcine ST and PK-15 cell lines, but rVSV-SADS did not replicate in these cells (data not shown). Therefore, further work is needed to conclude on the human specificity of the adaptive mutations we detected.

To understand the mechanistic basis of the increased fitness associated with SADS spike variants, we measured their processing and cell-cell fusogenicity. Interestingly, two mutations, V362M and L706G, significantly reduced SADS-CoV spike-mediated syncytia formation. In fitness assays, the V362M variant was neutral or deleterious in all cell lines, whereas the L706Q was beneficial in OVCAR-8 cells. Therefore, no direct link between SADS-CoV spike-mediated cell-cell fusion and viral fitness can be established. Many viruses, including SADS-CoV, induce cell-cell fusion during viral infection. Although it has been suggested that syncytia formation may facilitate viral replication, dissemination, pathogenesis or immune evasion (36), the link between cell-cell fusion and viral fitness is generally poorly understood. It has been shown that SADS-CoV spike cleavage at the FCS is necessary for efficient syncytia formation (35), in agreement with our data showing that the V362M mutation decreases both spike processing and cell-cell fusion. However, this mutation is located within the putative SADS-CoV spike receptor-binding domain (RBD), away from the FCS, and how it decreases spike cleavage remains unclear. Finally, the L706Q mutation decreased syncytia formation without altering spike processing, suggesting that factors other than spike cleavage are strong determinants of the SADS-CoV spike cell-cell fusogenicity.

Some of the mutations fixed during the experimental evolution clearly increased viral fitness but had no effect on spike cleavage or cell-cell fusion. These mutations may still represent adaptations to entry into human cells but may alter a process we have not looked at. For instance, an important aspect of viral entry that we could not study here is receptor binding. Indeed, to date, the SADS-CoV receptor remains unknown. It has been shown that SADS-CoV does not use other known coronavirus receptors (9), and although genome-wide CRISPR screens allowed the identification of host factors involved in SADS-CoV replication, they were unable to identify its receptor (37, 38). Interestingly, the A362M mutation is located with the S1 CTD, which usually serves as the coronavirus RBD for proteinaceous receptors. Once the receptor is identified, it will be interested to test whether this mutation has an impact on receptor affinity. Similarly, we have not looked at the effect of the identified mutations on the binding to other attachment factors (e.g. cell surface sugars) or processing by proteases (e.g. TMPRSS13) (19, 39). Moreover, although all mutations appeared together with others, we tested them all individually. Previous work on SARS-CoV-2, for example, has nonetheless shown that epistasis can strongly impact the effect of a spike mutation on receptor affinity or antibody escape (40, 41). Whether epistasis between the identified SADS-CoV spike mutations may alter their effect on viral fitness or spike processing and incorporation deserves further investigation.

Interestingly, most identified mutations occurred in the S2 subunit of the spike, especially near HR1 (A750G, and V754M) and HR2 (S1037G and T1107I). As none of these mutations impacted significantly cell-cell fusion and spike processing, how they increase viral fitness remains unclear. In the context of SARS-CoV-2, several S2 mutations have been fixed in the successive variants of concern (42), suggesting that they provide a fitness advantage to the virus. For SARS-CoV-2 and other coronaviruses, it has been shown that such S2 mutations can alter receptor affinity, fusogenicity, entry pathways or antibody sensitivity, through poorly understood mechanisms (43, 44). Future work is needed to understand how the SADS-CoV spike S2 mutations we have identified impact viral entry.

We used a recombinant VSV to study the evolution of the SADS-CoV spike. Although this is a practical and reliable system to study viral entry, some differences between VSV and SADS-CoV may have impacted the evolution of the spike. Therefore, we cannot exclude that some of the identified mutations may represent adaptations to the rVSV system rather than true human adaptations. One notable difference difference between VSV and SADS-CoV is their budding site.

VSV buds at the plasma membrane whereas coronavirus budding occurs in the endoplasmic reticulum (ER)-Golgi compartment, where an ER-retention motif in the cytoplasmic tail of the spike facilitates its accumulation (45). The cytoplasmic tail truncations (Q1120* and G1130*) we observed may therefore disrupt the spike retention motif, altering the spike cellular localization and redirecting it to the plasma membrane to favor its incorporation into budding VSV particles (46). Interestingly, a rVSV-SADS was recently engineered and, similarly to our observations, an 11-amino-acid truncation of the spike cytoplasmic tail was fixed after six passages in Huh7.5.1 cells and enhanced rVSV-SADS replication (47). Future work, using for example recombinant SADS-CoV (37), will help determining whether the spike mutations we identified are adaptations to human cells or to the rVSV system used.

In conclusion, we have used experimental evolution to identify potential human-adaptive mutations in the SADS-CoV spike that may enhance viral entry into human cells. Further work aimed at understanding how SADS-CoV can adapt to different host species will be crucial to monitor SADS-CoV evolution in nature and prevent potential cross-species transmission events.

## Methods

### Cell lines

The NCI-60 panel (dtp.cancer.gov/discovery_development/nci-60) was purchased from the National Cancer Institute. Twelve cell lines showing poor growth, lack of susceptibility to VSV or poor adherence were excluded. Information about the remaining 48 cell lines is provided in **Supplementary Table 1**. OVCAR-8, H23, and other NCI-60 cells were cultured in RPMI supplemented with 10% FBS, 1% non-essential amino acids, penicillin (10 U/mL), streptomycin (10 µg/mL) and amphotericin B (250 ng/mL). Huh-7 cells were kindly provided by Ralf Bartenschlager (Heidelberg University Hospital). BHK-21 were obtained from the ATCC (ATCC, CCL-10). BHK-G43 cells, which can induce the VSV-G protein expression after mifepristone treatment (48), were kindly provided by Gr. Gert Zimmer. HEK-293T-GFP1-10 and HEK-293T-GFP-11 were both kindly provided by Olivier Schwartz (Institut Pasteur, France) and maintained in the presence of 1 µg/mL puromycin. Huh-7, BHK-21, BHK-G43, and HEK-293T cells were cultured in Dulbecco’s Modified Eagle’s Medium supplemented with 10% FBS, 1% non-essential amino acids, penicillin (10 U/mL), streptomycin (10 µg/mL) and amphotericin B (250 ng/mL). All cells were maintained at 5% CO_2_ and 37°C in a humidified incubator and were routinely screened for the presence of mycoplasma by PCR.

### Recombinant viruses

The rVSV bearing the SADS-CoV spike was generated using a previously described system (49). Briefly, the codon-optimised SADS-S (GenBank AVM80475.1) was ordered as a synthetic gene (GenScript) and cloned into a plasmid encoding the Indiana serotype VSV antigenome replacing the VSV glycoprotein gene. The plasmid was previously modified to encode eGFP from an additional transcription unit between the G and L genes (pVSV-eGFP-ΔG). BHK-G43 cells were seeded at a density of 10^5^ cells/mL in DMEM with 10% FBS and without antibiotics in 12-well plates (1 mL/well). On the day of transfection, the viral genome pVSV-eGFP-ΔG-SADS-S (25 fmol) was co-transfected with helper plasmids encoding VSV-P (25 fmol), VSV-N (75 fmol) and VSV-L (25 fmol) proteins and a codon-optimised T7 polymerase plasmid (50 fmol). Lipofectamine 3000 (Invitrogen) was used for transfection. Cells were incubated with the transfection mix at 37°C for 3 hours. Then, 1 mL of DMEM 10% FBS supplemented with 10 nM mifepristone was added to each well and the cells were incubated at 33°C for 36 h, followed by 36-48 h at 37°C. The supernatant collected from GFP-positive cells was purified by centrifugation at 2000 g for 10 min and used to infect VSV-G-induced BHK-G43 (seeded the day before at 10^5^ cells/mL in DMEM 10% FBS in 12-well plates) for 1 h at 37°C, in order to increase viral titers. The supernatant was harvested at 36 h post-inoculation (hpi), cleared by centrifugation at 2000 g for 10 min and aliquoted. To obtain viruses without VSV-G, BHK-21 cells were infected with the supernatant from previous amplification in BHK-G43. Following 1 h at 37°C, the inoculum was removed, the cells were washed five times with PBS, and incubated in complete DMEM supplemented with 25% anti-VSV-G neutralizing monoclonal antibody obtained in-house from a mouse hybridoma cell line. The supernatant was collected at 36 hpi, cleared by centrifugation at 2000 g for 10 min, aliquoted, and stored at -80°C.

### rVSV-SADS experimental evolution

The SADS-CoV S-expressing recombinant VSV was serially passaged in triplicate in Huh-7, H23 and OVCAR-8 cell lines until 10 passages were completed. For each passage, 6-well confluent plates were infected with 200 µL of virus at a multiplicity of infection (MOI) of ≤ 0.01 FFU/cells and incubated for 1 h at 37°C and 5% CO_2_, with agitation every 15 min. Then, 2 mL of complete DMEM with 2% FBS was added to each well. After 3-5 days at 37°C and 5% CO_2_ (3 days in Huh-7, 5 days in H23 and OVCAR-8), the supernatants were collected, clarified by centrifugation at 2000 g for 10 min, aliquoted and stored at -80°C. The harvested supernatants were then used to infect new 6-well plates with confluent monolayers, thus initiating the subsequent passage. Supernatants were titrated between each passage, as detailed below.

### Virus titration

Supernatants were serially diluted and 100 µL were used to infect 24-well confluent Huh-7 plates for 1 h at 37ºC. 500 µL of DMEM with 2% FBS and 2% agar was then added to each well. The plates were incubated for 18-24 h at 37°C and 5% CO_2_. The plates were imaged in the Incucyte SX5 Live-Cell Analysis System (Sartorius) in order to manually count the GFP-positive foci. Viral titers were expressed as FFU/mL.

### Sanger sequencing

Viral RNA was extracted using the NZY Viral RNA Isolation kit (NZYtech, MB40701) following the manufacturer’s instructions. The extracted RNA was reverse transcribed (RT) using SuperScript IV (Invitrogen) with a VSV backbone-specific primer (5’-CTCGAACAACTAATATCCTGTC-3’). The cDNA was purified using the DNA Clean & Concentrator-5 kit (Zymo Research) and used for the spike gene amplification with Phusion Hot Start II High-Fidelity PCR Master Mix (ThermoFisher) using VSV backbone-specific primers (5’-CTCGAACAACTAATATCCTGTC-3’, 5’-GTTCTTACTATCCCACATCGAG-3’). The PCR product was then sent to Sanger sequencing using the VSV-specific primers previously mentioned and spike-specific primers listed in **Supplementary Table 3**.

### Site-directed mutagenesis

SADS-CoV spike mutations were inserted into the SADS glycoprotein encoding plasmid template (pcDNA3.1-SADS-S-HisTag) by site-directed mutagenesis using the QuickChange II XL Site-Directed Mutagenesis Kit (Agilent, 200522) according to the manufacturer’s instructions. Each reaction was made with 30 ng of template, dNTPs (25 nM each), 250 ng of each primer pair and 2.5 U/µL of PfuUltra HF DNA polymerase. The primers used are listed in **Supplementary Table 4**. SDM products were digested at 37ºC for 1h with FastDigest DpnI (Thermo Scientific) and transformed into NZY5α competent cells (NZYTech). The presence of the desired mutation in specific clones was confirmed by Sanger-sequencing. Plasmids with the desired spike mutations were then used for PCR using Phusion Hot Start II High-Fidelity PCR Master Mix (ThermoFisher). A pair of primers was used to amplify each spike (5’-CGATCTGTTTACGCGTCACTATGAAGCTGTTCACCGTG-3’; 5’-AGCAGGATTTGAGTTAATCGTTAATGGTGATGGTGATGG-3’). The PCR product was purified using the DNA Clean & Concentrator-5 kit (Zymo Research) and cloned with HiFi (NEBuilder) into the pVSV-eGFP-ΔG plasmid previously described, using a pair of primers to open the backbone (5’-CGATTAACTCAAATCCTGC-3’; 5’-AGTGACGCGTAAACAGATC-3’). A whole plasmid sequencing was done to confirm the presence of the desired mutation.

### Cell-cell fusion assay

The cell-cell fusion assay was performed as previously described (50). HEK293T-GFP1-10 and HEK293T-GFP11 were mixed at a 1:1 ratio and 6 ×10^5^ cells per well in a 96-well plate were transfected in suspension with 100 ng of pcDNA3.1-SADS-spike plasmid (WT or mutants) or an empty plasmid as a control using Lipofectamine 2000 (InVitrogen). The plates were placed in an Incucyte SX5 Live-Cell Analysis System (Sartorius) at 37°C and 5% CO_2_ for imaging. The percentage of fusion was calculated as the ratio of the GFP area to the cell confluence area at 18-21 h post-transfection, as measured using the Incucyte analysis software.

### Quantitation of viral spread and endpoint titers

Huh-7, OVCAR-8 and H23 cells were plated at 60% confluence in 12-well plates. The following day, cells were inoculated with 100 µL of virus dilution at a MOI of approximately 0.004 FFU/cell. Cells were incubated at 37°C with 5% CO_2_ and agitated every 15 min. Following 1.5 h, 1 mL of DMEM containing 2% FBS was added, and cells were incubated for 3 days (Huh-7) or 5 days (OVCAR-8 and H23) at 37°C and 5% CO_2_. The plates were imaged at 6 h intervals using the Incucyte SX5 Live-Cell Analysis System (Sartorius) to determine the area occupied by GFP-positive cells. The end-point supernatants were harvested, clarified by centrifugation (2000 g for 10 min), and stored at -80°C. Viral titers were determined as described above.

### Western blotting

Supernatant containing recombinant viruses was centrifuged at 30,000 g for 2 h at 4°C. Viral pellets were lysed in 30 µL of NP-40 lysis buffer (Invitrogen) supplemented with a complete protease inhibitor (Roche) for 30 min on ice. Lysates were mixed with 4x Laemmli buffer (Bio-Rad) supplemented with 10% β-mercaptoethanol and denatured at 95°C for 5 minutes. Proteins were separated by SDS-PAGE using pre-cast 4-20% Mini-PROTEAN TGX Gels (Bio-Rad) and transferred onto a 0.45 µm PVDF membrane (Thermo Scientific). Membranes were blocked with TBS-T (20 mM tris, 150 nM NaCl, 0.1% Tween-20, pH 7.5) supplemented with 3% bovine serum albumin for 1 h at room temperature. The membranes were then incubated for 1 h at room temperature with two primary antibodies: mouse anti-6X-HisTaq (dilution 1:1.000, MA1-21315, Invitrogen) and mouse anti-VSV-M (dilution 1:1.000, clone 23H12, EB0011, Kerafast). Following three washes with TBS-T, the primary antibodies were detected using a goat anti-mouse IgG secondary antibody conjugated to horseradish peroxidase (HRP) (dilution 1:50,000, G21040, Invitrogen). The signal was revealed using PierceTM ECL Plus Western Blotting Substrate (32132, Thermo Scientific), following the manufacturer’s instructions. Images were captured using an ImageQuant LAS 500 (GE Healthcare) and analysed using Fiji software (v.2.14.0).

### Statistics

All statistical analyses were conducted using GraphPad Prism v10.2.3. Details on the statistical tests employed are provided in the main text and in the figure legends.

## Acknowledgments

We thank all members of the laboratory for helpful discussions about this work. We thank Iván Andreu-Moreno and Raquel Martínez-Recio for help in the generation of the rVSV-SADS WT. This work was financially supported by ERC Advanced Grant 101019724-EVADER, grant PID2020-118602RB-I00 from the Spanish Ministerio de Ciencia e Innovación (MICINN), and grant CIAICO/2022/110 from the Conselleria de Educación, Universidades y Empleo (Generalitat Valenciana) to R.S. J.D. was supported by an EMBO postdoctoral fellowship (ALTF 140-2021) and a Marie Skłodowska-Curie Actions Postdoctoral Fellowship (101104880).

## Author contributions

R.S. and J.D. designed research; C.S.-T performed research; C.S.-T., R.S. and J.D. analyzed data; C.S.-T, J.D. and R.S. wrote the paper; R.S. provided funding.

